# Population Genotype Calling from Low-coverage Sequencing Data

**DOI:** 10.1101/085936

**Authors:** Lin Huang, Petr Danecek, Sivan Bercovici, Serafim Batzoglou

## Abstract

In recent years, several large-scale whole-genome projects sequencing tens of thousands of individuals were completed, with larger studies are underway. These projects aim to provide high-quality genotypes for a large number of whole genomes in a cost-efficient manner, by sequencing each genome at low coverage and subsequently identifying alleles jointly in the entire cohort. Here we present Ref-Reveel, a novel method for large-scale population genotyping. We show that Ref-Reveel provides genotyping at a higher accuracy and higher efficiency in comparison to existing methods by applying our method to one of the largest whole-genome sequencing datasets presently available to the public. We further show that utilizing the resulting genotype panel as references, through the Ref-Reveel framework, greatly improves the ability to call genotypes accurately on newly sequenced genomes. In addition, we present a Ref-Reveel pipeline that is applicable for genotyping of very small datasets. In summary, Ref-Reveel is an accurate, scalable and applicable method for a wide range of genotyping scenarios, and will greatly improves the quality of calling genomic alterations in current and future large-scale sequencing projects.

---

Several large-scale whole-genome sequencing projects have been completed, while even more extensive efforts are currently underway (The 1000 Genomes Project Consortium, 2010; The UK10K Consortium, 2015; CHARGE Consortium, 2009; CONVERGE consortium, 2015; <u>http://www.haplotype-reference-consortium.org</u>; <u>https://www.genomicsengland.co.uk/</u>). These projects were designed to characterize human genetic variation across various populations, enabling subsequent association studies that aim to identify the underlying mechanisms that drive human hereditary disease. For each one of these studies, once the large cohorts are defined, collected, and sequenced, genotype calling is applied on a massive volume of whole genome sequencing data. The accuracy at which the genomic variation is identified strongly influences the quality and statistical power of downstream analyses. Given the size of such projects, conducted in both academia and industry, there is a growing need for the ability to infer the genotypes across a very large sequencing datasets (4,000 – 1,000,000 whole genomes). As the size of the data size increases, traditional population genotype calling methods become prohibitively slow. As such, an accurate and scalable computational method for population genotype calling is urgently needed.

To characterize human genetic variation, significant research efforts and massive resources have been expended in these projects to sequence a large number of whole genomes and to call genotypes of the sequenced genomes at polymorphic sites. Individual-level genomic data from these projects is available to the scientific community. These resources, beyond their value in the original projects and genome-wide association studies, can be used to enhance the quality of population genotyping in future genome sequencing projects. The genotypes identified capture the linkage disequilibrium (LD) structure of multiple populations. Cohorts that will be sequenced in future projects are likely to share a similar LD structure with one or more studied cohorts. Given the fact that this insight is being leveraged today in many haplotyping and imputation methods (Delaneau et al., 2014; Huang et al., 2015), the genotype calls from completed projects can be used as a reference panel in future genotype calling process.

While advancements in sequencing technology have enabled a sharp reduction in sequencing cost, the cost becomes prohibitive again once hundreds of thousands, or even millions, of individuals are targeted for sequencing. In such scenarios, low-coverage whole genome sequencing of a large cohort provides a promising path forward, enabling the efficient identification of genomic alterations. Many recent sequencing projects, including the ones mentioned above, have employed the low-coverage/large-dataset strategy. For instance, the 1000 Genomes Project sequenced 2,504 whole genomes at a mean depth of 7.4x; the UK10K Cohorts Project sequenced 3,781 whole genomes at a mean read depth of 7x. This strategy inevitably induces the loss of information at every single site, which is one of the main difficulties in the process of genotype calling. Without incorporating prior information from additional sources, the coverage requirement of population sequencing projects can hardly be further reduced. Reference-guided genotype calling, which incorporates such priors, has the potential to enable a low-coverage sequencing without sacrificing genotype calling accuracy. In other words, larger studies can be supported, sequencing additional individuals, and enabling an even broader investigation of the underlying genetic makeup.

A number of population genotyping methods have been proposed to call genotypes from large-scale low-coverage whole genome sequencing data. The examples include glfMultiples+Thunder (Li et al., 2011), which employs a hidden Markov model that leverages LD information across a cohort to genotype likely polymorphic sites; SNPTools (Wang et al., 2013), which estimates genotyping likelihoods using a BAM-specific binomial mixture model and then utilizes a hidden Markov model (HMM) approach based on the statistical LD pattern model proposed in (Li and Stephens, 2003) to infer genotypes and haplotypes; and Beagle (Browning and Yu, 2009), which builds a HMM-based haplotype frequency model to capture LD pattern. These methods analyze all the sequenced samples in a cohort jointly, because calling genotypes from the data of a single low-coverage sequenced sample yields poor results. Despite the considerable success of these methods, using HMM-based models inevitably involves undesirable scalability and tendency to weaken long-distance LD, which is critical for calling the genotypes of rare variants (Huang et al., 2016). Both issues make these methods less suitable for large-scale population genotyping.

To address these issues, we present reference-based Reveel (or Ref-Reveel), a novel population genotyping method. Compared to previous methods, Ref-Reveel provides higher genotyping accuracy across the allele frequency spectrum, especially at rare variants, while maintaining a low computational overhead. Additionally, when a cohort of genotyped reference individuals is available, Ref-Reveel can leverage these genotypes as a *reference panel* to accurately infer the genotypes of a newly low-coverage sequenced cohort of individuals, referred to as *query genomes*.

Ref-Reveel provides significant enhancements over our previous methods, Reveel (Huang et al. 2016). In our new method, Ref-Reveel effectively incorporates genotypes from completed projects, if applicable, to improve the genotyping quality of new datasets while maintaining low computational costs. Ref-Reveel creates a family of weak learners that exhibit low computational cost, and learns the promising combination through an AdaBoost-based (Freund and Schapire, 1997) meta-algorithm for every marker from the newly sequenced genomes. Furthermore, by incorporating a reference panel, Ref-Reveel achieves high genotyping quality when applied very small datasets. Finally, several computational challenges are resolved to enable the application of Ref-Reveel on thousands to tens of thousands of whole genomes, simultaneously, resulting in the producing of accurate calls within a reasonable time-frame.

## Results

### Genotype calling the whole genomes of 3,910 UK10K samples

We first show that Ref-Reveel is a practical infrastructure for inferring the genotypes of a large number of whole genomes. When applied to one of the largest data sets that are currently publicly available, Ref-Reveel outperformed state-of-the-art methods, both in terms of genotyping quality as well as efficiency. In this experiment, we applied Ref-Reveel to the genotype likelihoods of 3,910 UK10K samples from two British cohorts, the Avon Longitudinal Study of Parents and Children and TwinsUK, to infer the genotypes of these samples. This input dataset included 61,897,468 unfiltered sites across the whole genome. The input genotype likelihoods were calculated from low-coverage sequencing data (average read depth 7x) using SAMtools and BCFtools by the UK10K Consortium.

We performed this experiment in parallel on Sanger Institute’s computing farm. To do so, we first partitioned the whole genome into 4,720 non-overlapped genomic segments. The chromosomes were chunked by marker numbers; the maximum number of markers per segment was set to 12,000. Then, Ref-Reveel was applied to each segment separately. A total of 61,167,575 sites were genotyped by Ref-Reveel across the entire genome; the other 729,893 sites were identified by Ref-Reveel as invariant reference alleles across the studied cohort. Ref-Reveel produced genotype likelihoods at common and low-frequency variation sites and genotypes at rare and invariant sites. The genotype likelihoods at common and low-frequency variation sites, which were roughly 13% of all the genotyped sites, were fed into Beagle (version 3.3.2) for the final refinement. The refined genotypes were merged with the genotypes at rare and invariant sites. The total running time for the combined pipeline, when applied on the above dataset, was 85,211 CPU hours, out of which Ref-Reveel consumed 49,856 CPU hours (59% of the total running time); the memory usage was 18.6 GB.

We compared the panel (labelled as *R*) with the haplotypes of 3,781 UK10K samples reported by the UK10K Consortium (labelled as *10K*). The 10K panel was created as follows by the UK10K Consortium (The UK10K Consortium, 2015). The SNP calls in panel 10K were made using SAMtools and BCFtools, then recalled using the GATK (version 1.3-21) UnifiedGenotyper, and filtered using the GATK VariantRecalibrator followed by GATK ApplyRecalibration. The missing and low confidence genotypes were then refined using Beagle 4 (rev909). The UK10K Consortium reported 42,001,233 polymorphic sites that passed the filter; the haplotypes at these sites were included in the panel.

We measured the quality of these two reference panels on 66 TwinsUK samples that were both low-coverage whole-genome sequenced and high read-depth exome sequenced. The genotype calls from high read-depth exome sequencing reported by TwinsUK were used as benchmarks, including 160,119 sites. We evaluated genotyping accuracy, sensitivity and precision as defined in **Table 1** at the polymorphic sites discovered from the exome sequencing data.

**Table 1.**
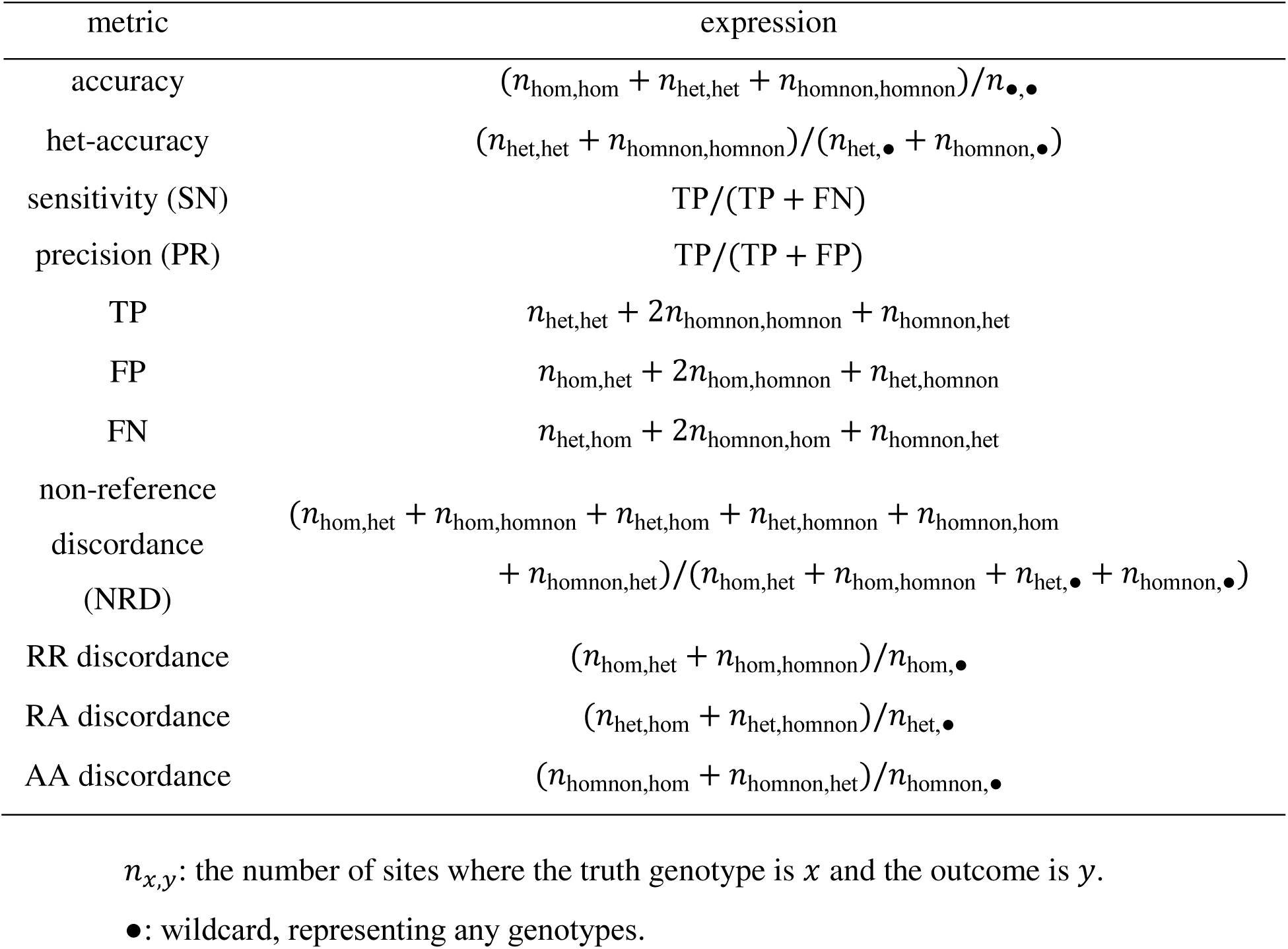
Metrics used for genotyping performance assessment.

**Figure 1** clearly shows that panel R achieved higher quality in comparison to panel 10K. A closer examination shows that panel R had moderately lower precision and significantly higher sensitivity than panel 10K. The result was expected, because when Ref-Reveel was used to generate a reference panel, its SNP discovery parameter was set to a very low value to infer as many likely variation sites as possible. Despite its higher false positive rate, we used this parameter setting as our experiments show that a panel with high SNP discovery rate benefited the downstream reference-based genotype calling (see **Incorporating reference panels to facilitate genotype calling of CEU samples**).

**Figure 1.**
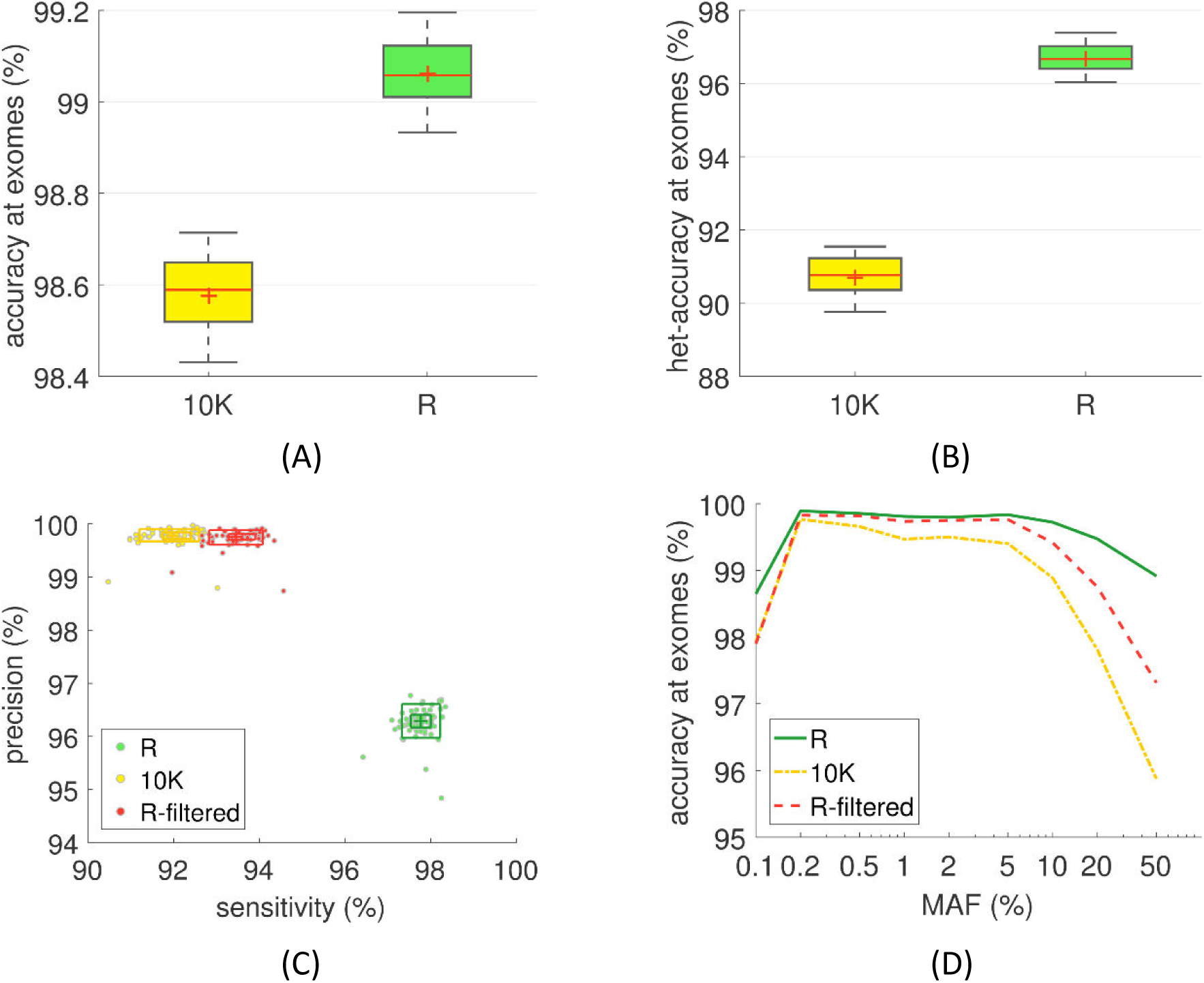
Quality of two UK10K reference panels. We showed the aggregated (A) accuracy and (B) het-accuracy of 66 TwinsUK samples using boxplots, in which the central marks are the mean, the red lines are the median, the edges of the box are the 25th and 75th percentiles, and the whiskers span 9th to 91st percentiles. We also showed the sensitivity and precision using rangefinder boxplots (C). The dots are the performance of individuals; the inner and outer boxes span the 25th-75th and the 9th-91th percentiles respectively. The average accuracy across the whole genome as a function of minor allele frequency is shown in (D).

To compare the Ref-Reveel-called panel and panel 10K at the same precision level, we applied four hard filters described in **Online Methods** to reduce the false positive rate of panel R. Across the whole genome 2,310,217 sites (3.78%) were removed by these filters. As shown in **Figure 1(C)**, the sensitivity of the filtered Ref-Reveel-called panel was higher than that of panel 10K.

The superiority of panel R over panel 10K was observed across the whole allele frequency (AF) spectrum, but the advantage at the sites with minor allele frequency (MAF) <0.1% and MAF≥1% was particularly significant (**Figure 1(D)**). For rare variants, Ref-Reveel had higher genotyping accuracy because of its unique capability of identifying the most informative sites in a way that is less sensitive to genetic distance. Other methods, by contrast, implicitly weaken the association between remote sites when they build their models. For high AF variants, we attributed the high genotyping accuracy of Ref-Reveel to its another important feature, that is, Ref-Reveel provided high-quality genotype probabilities at high AF sites. Applying the hard filters to panel R cancelled the advantage at the very rare sites, implying that a good amount of very rare variants was removed by the hard filters. This could weaken the power of panel R as a reference panel. Applying the hard filters also reduced the accuracy at common variants. Even though, the filtered R panel still had higher quality than panel 10K at those sites.

### Incorporating reference panels to facilitate genotype calling of CEU samples

To demonstrate the improved performance of Ref-Reveel gained through the incorporation of a reference panel, we applied our method on real low-coverage data, contrasting our calls against an orthogonal validation set. In this experiment, we applied Ref-Reveel on 99 CEU samples from the 1000GP Phase 3. The BAM files corresponding to the low-coverage sequencing data were retrieved from the 1000GP website (retrieved on 01/13/2016). Sequentially, we applied SAMtools (version 1.2)’ mpileup command to generate the genotype likelihoods. We used the application of reference-free Ref-Reveel as the baseline for our comparison, denoted as *R-baseline*.

We assessed the performance of Ref-Reveel using three reference panels. The first two reference panels were panel 10K and panel R, as described above. The third reference panel, denoted as 1000GP, contained the haplotypes of 2,405 non-CEU 1000GP samples from the integrated call set reported by the 1000GP Phase 3 (ftp://ftp.1000genomes.ebi.ac.uk/vol1/ftp/release/20130502/). The resulting genotype calls using these three reference panels were labelled as *R-10K*, *R-R*, and *R-1000GP*, respectively.

We compared Ref-Reveel against three state-of-the-art pipelines. In the first pipeline, we combined SNPTools (Wang et al., 2013) and Beagle (Browning and Browning, 2009), denoted as *S+B*. To estimate the genotype likelihoods at polymorphic sites, we used SNPTools (v1.0)’ bamodel→poprob commands. We then applied Beagle 4 (r1399) to infer the final set of genotypes. For the second pipeline, we combined GATK (DePristo et al., 2011; McKenna et al., 2010) with Beagle, denoted as *G+B*. Namely, Beagle 4 (r1399) used the genotype likelihoods generated by GATK UnifiedGenotyper (v3.3) to infer the final set of genotypes. Finally, our third pipeline was glfMultiples+Thunder (Li et al., 2011), denoted as *g+T*. While Thunder is computationally intensive, it has demonstrated improved genotyping accuracy when applied to the output of glfMultiples. All methods were applied using default parameters, unless otherwise specified.

To assess performance, we contrasted the generated calls against the genotypes reported in the Complete Genomics (CG) dataset (retrieved on 01/13/2016). A total of 63 samples, out of the initial 99 samples described above, were reported in the CG dataset. We measured genotyping accuracy at all the sites where the CG data reported heterozygous and homozygous alternate, that is, het-accuracy defined in **Table 1**.

The performance of R-baseline, R-10K, R-R, and R-1000GP as evaluated across the entire human genome is outlined in **Table 2**. We focused on the sites that were both reported by the CG data and genotyped by Ref-Reveel. Besides measuring the overall performance, we further divided the sites into four categories: common variants, denoted as *S*_common_; low frequency variants, denoted as *S*_loF_; rare variants, denoted as *S*_rare_; and finally, invariant sites, denoted as *S*_invariant_, for sites that do not appear in the integrated call set of the 1000GP Phase 3. The performance was evaluated for each variant category separately.

**Table 2.** Genotyping accuracy of Ref-Reveel on the whole genome. The genotyping accuracy was evaluated at the sites where the CG data reported heterozygous and homozygous alternate and Ref-Reveel discovered SNPs.

| method | overall (%) | $S_{\text{common}}$ (%) | $S_{\text{loF}}$ (%) | $S_{\text{rare}}$ (%) | $S_{\text{invariant}}$ (%) |
| --- | --- | --- | --- | --- | --- |
| R-baseline | 98.7267 | 99.1241 | 98.4575 | 97.4269 | 80.5800 |
| R-10K | 98.8818 | 99.2576 | 98.8222 | 97.9519 | 79.7389 |
| R-R | 99.0280 | 99.3671 | 99.0418 | 98.1839 | 81.4276 |
| R-1000GP | 98.9069 | 99.3089 | 98.7196 | 97.7574 | 79.5758 |

Given the above dataset, Ref-Reveel discovered 25,261,018 likely polymorphic sites, and reported invariant alleles at remaining sites. A general trend shows that using a reference panel improved the performance across the entire allele frequency spectrum compared to R-baseline. Compared to the baseline, using the UK10K samples as references improved the genotyping accuracy. The observed improvement in genotyping accuracy and sensitivity was not surprising since the British cohort and the CEU samples exhibited low genetic divergence (Eyheramendy et al., 2015; The 1000 Genomes Project Consortium, 2015). As such, we expected that indeed the UK10K reference panel would provide a proper approximation of the LD structure in the CEU population. Notably, R-R exhibited a higher genotyping accuracy in comparison to the R-10K, although both reference panels originated from the UK10K dataset. We attributed the observed result to the improved genotyping quality of panel R. At the common variants, using panel R reduced the genotyping error rate from 0.88% to 0.63%, corresponding to the genotypes of 7,847 markers per sample corrected; at the low-frequency variants, we observed the reduction from 1.54% to 0.96%, corresponding to the genotypes at 1,089 markers per sample corrected; at the rare variants, the error rate was reduced from 2.57% to 1.82%, corresponding to the genotypes at 878 markers per sample corrected. It is important to note that while the 1000GP reference panel originated from a more heterogeneous set of reference populations, when that panel was utilized for genotype calling, compared to the baseline we observed a 21.1%, 17.0%, 12.8% reduction in the genotyping error rate at common, low-frequency, and rare variants, respectively. This could imply that the LD structure of the variants with sufficiently high AF were likely to be captured by this reference panel, and that with the AF decrease the LD structure became less likely to be captured.

We contrasted the performance of Ref-Reveel against state-of-the-art methods over chromosome 20, which was roughly 2% of the whole genome; assessing the performance of the alternative methods across the entire genome was computational prohibitive. The performance was evaluated at all the sites where the CG data reported either heterozygous or homozygous alternate on this chromosome. As shown in **Figure 2**, Ref-Reveel, with or without a reference panel, outperformed previous state-of-the-art methods. The SNPTools+Beagle and GATK+Beagle pipelines performed worse than any setting of Ref-Reveel in terms of het-accuracy. The third pipeline, glfMultiples+Thunder, achieved higher accuracy in comparison to the two previous ones, yet exhibited a lower performance in comparison to the R-baseline results.

**Figure 2.**
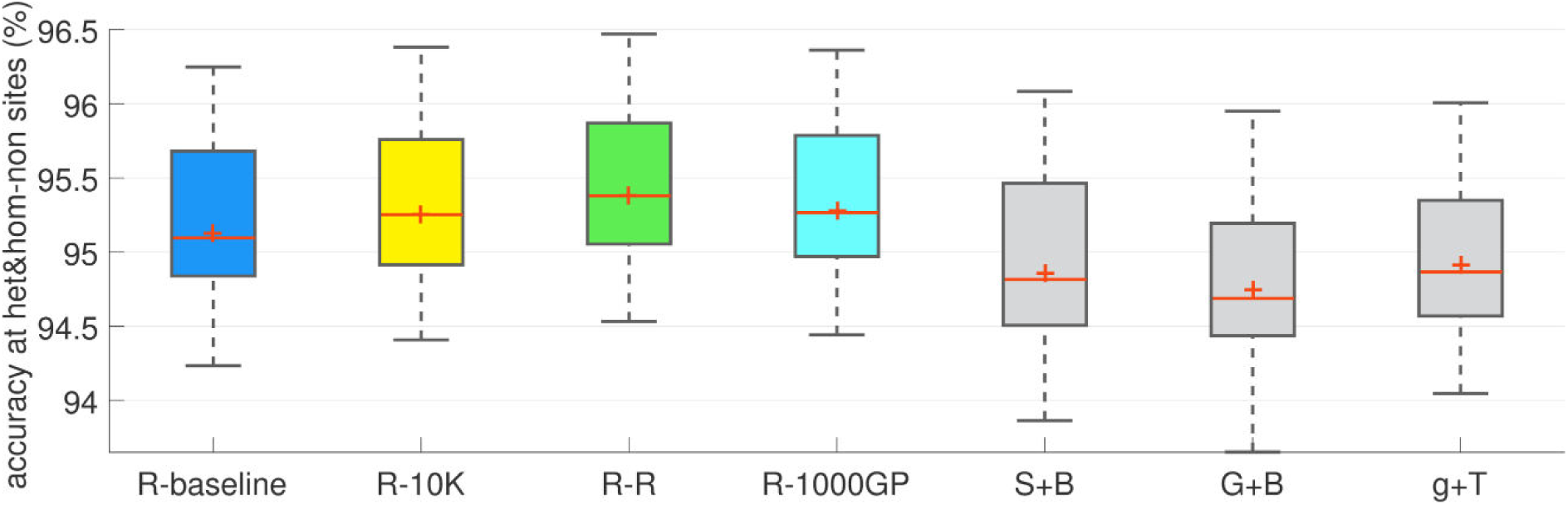
Genotyping performance for different methods and reference panels on chromosome 20. We compared four settings of Ref-Reveel with different reference panels, labelled as R-baseline, R-10K, R-R, and R-1000GP, against three state-of-the-art pipelines, labelled as S+B, G+B, and g+T, on 99 CEU samples from the 1000GP Phase 3. For each method or setting, the genotyping accuracy at heterozygous and homozygous alternate sites of 63 samples that were studied by both 1000GP Phase 3 and CG was represented by a boxplot.

As shown in **Table 3**, R-baseline had the lowest CPU running time of 7.36 CPU days when evaluating calls across the entire genome, and a total of 134 minutes when evaluating calls on chromosome 20 alone. Incorporating the reference panels roughly doubled the running time. When panel R was used, Ref-Reveel’s computational time was 12.79 CPU days on the whole genome, and 353 minutes on chromosome 20. This computational overhead is practical, even when a single core is used. Given the fact that Ref-Reveel is parallelizable in a straightforward manner (by analyzing independent genomic regions in parallel). A cluster can be used to reduce the end-to-end running time to less than a day.

**Table 3.** Running time. The running time was measured on a 2.40GHz Intel Xeon processor.

| method | chromosome 20 (mins) | whole genome (CPU days) |
| --- | --- | --- |
| R-baseline | 134 | 7.36 |
| R-10K | 323 | 13.02 |
| R-R | 353 | 12.79 |
| R-1000GP | 275 | 11.55 |
| S+B | 407 | - |
| G+B | 1464 | - |
| g+T | 21924 | - |

Among the state-of-the-art methods, SNPTools+Beagle was the only one that had efficiency comparable with that of Ref-Reveel. According to our experiment on chromosome 20, GATK+Beagle was 4.2 times slower than R-R, whereas glfMultiples+Thunder was 62.1 times slower. Extrapolating from these results, one can estimate that the running time of these two pipelines on the entire genome will be approximately 53 CPU days and 794 CPU days respectively. We conclude that Ref-Reveel provides substantial accuracy and efficiency improvements in population genotyping, and enables the accurate and efficient genotyping of a sequenced cohort using a previously genotyped reference panel cohort.

We also conducted a simulation study to demonstrate the effectiveness of incorporating reference panels on a larger query sample size (**Supplementary Note 1**, **Supplementary Figure S1** and **Supplementary Table S1**). The genotyping quality was greatly improved across the allele frequency spectrum for all three cases in which the query sample sizes were 100, 500, and 1000. In the meantime, although parsing the reference VCF file and incorporating the reference haplotypes into the genotyping process introduced additional computation overhead, the additional running time scaled well with the increase of sample size.

### Ref-Reveel powers high-accuracy calls on small datasets

Utilizing reference panels enables the genotype calling of a small (<50 samples), low-coverage dataset, which is a challenging scenario for all the population-genotyping tools we have reviewed including Reveel. Genotype calling from low-coverage sequencing data highly relies on the ability of genotype caller to discover the underlying complex LD structures. A small query dataset will cause inaccurate or even misleading estimation of linkage disequilibrium based on limited data. Incorporating a reference panel is a promising way to overcome this difficulty.

When the query samples come from heterogeneous, unknown populations, the scenario becomes even more challenging because picking a reference panel that matches the query samples is not straightforward. A poor representation of the query cohort by the reference panel will hamper the ability of genotype callers to leverage the underlying complex LD structure in the datasets. Ideally, the reference panel is composed of samples from the same population or ancestry group as the query samples.

To demonstrate the performance of the Ref-Reveel pipeline (**Online Methods**) to very small datasets, we conducted experiments as follows. We created the query dataset by randomly selecting ten samples from the 1000GP samples. These ten samples come from ten distinct populations from Africa, East Asia, Europe, and South Asia. Only the genotype likelihoods, generated using SAMtools, of these samples were included in the query dataset. The integrated call set reported by the 1000GP Phase 3, excluding the genotypes of these ten samples, was used as the reference panel. The resulting reference panel contained the genotypes of 2,494 samples, representing the original 26 populations. We evaluated the performance on a 5-Mbp region on chromosome 20 (43,000,000-48,000,000). In the experiment, for every query sample we picked five populations that are most similar with the query sample based on *F*-statistics and combined the 1000GP samples from these populations with that sample. This step gives a reasonable number of samples in the combined VCF file, ranging from 462 to 515. The running time of the first step was 3.65 minutes on a 2.40GHz Intel Xeon processor. The second step had negligible running time. Finally, the third step run time was 2.02 hours, per run.

We measured the genotyping performance of these query samples using the Omni genotype data as benchmarks. The performance was evaluated at 4,359 overlapped polymorphic sites. Beyond the estimated genotyping accuracy, we further measured the non-reference discordance (NRD), RR discordance, RA discordance, and AA discordance, as defined in **Table 1**. As shown in **Table 4**, the pipeline achieved a superior performance in all the query samples. Except for NA19430 (sample 9), the genotyping accuracy of all the samples ranged between and 99.93%. Out of 10 query samples, 4 samples had NRD < 1%, 5 samples had NRD 1-2%, and only a single sample, NA19430, had an NRD 3.14%. Examination of these numbers indicated that the sequencing depth strongly affected the genotyping performance. To support this hypothesis, we downsampled the BAM files of the query samples to 83.3%, 66.7%, 50%, and 33.3% of the original coverage level, recalculated the genotype likelihoods using SAMtools, and repeated the analysis above. The results for the three sets of query samples, each containing 10 query samples, are illustrated in **Figure 3**. Each gray curve in the plot corresponds to the performance of a query sample, whereas each colored curve corresponds to an aggregated performance of the samples originating from a single continent. The figure clearly shows the correlation between the sequencing depth and the genotyping performance. The populations from which the samples originated, on the other hand, was not a critical factor. As shown in **Figure 3**, the aggregated curve of four continents nearly overlaps. The evaluation can be used to provide guidance when one wishes to initiate a sequencing project focusing on a new cohort. When we call genotypes of a very small set of query samples using the above described pipeline, a sequencing depth of 4x or higher is expected to result in a sample’s genotyping accuracy of >99%; a sequencing depth of 2x is sufficient to achieve a mean genotyping accuracy of 99% across a cohort. Finally, a sequencing depth of 8x or higher will often reduce the NRD below 1%. When the pipeline is applied to a larger query dataset, we expect the coverage requirement to be further reduced.

**Figure 3.**
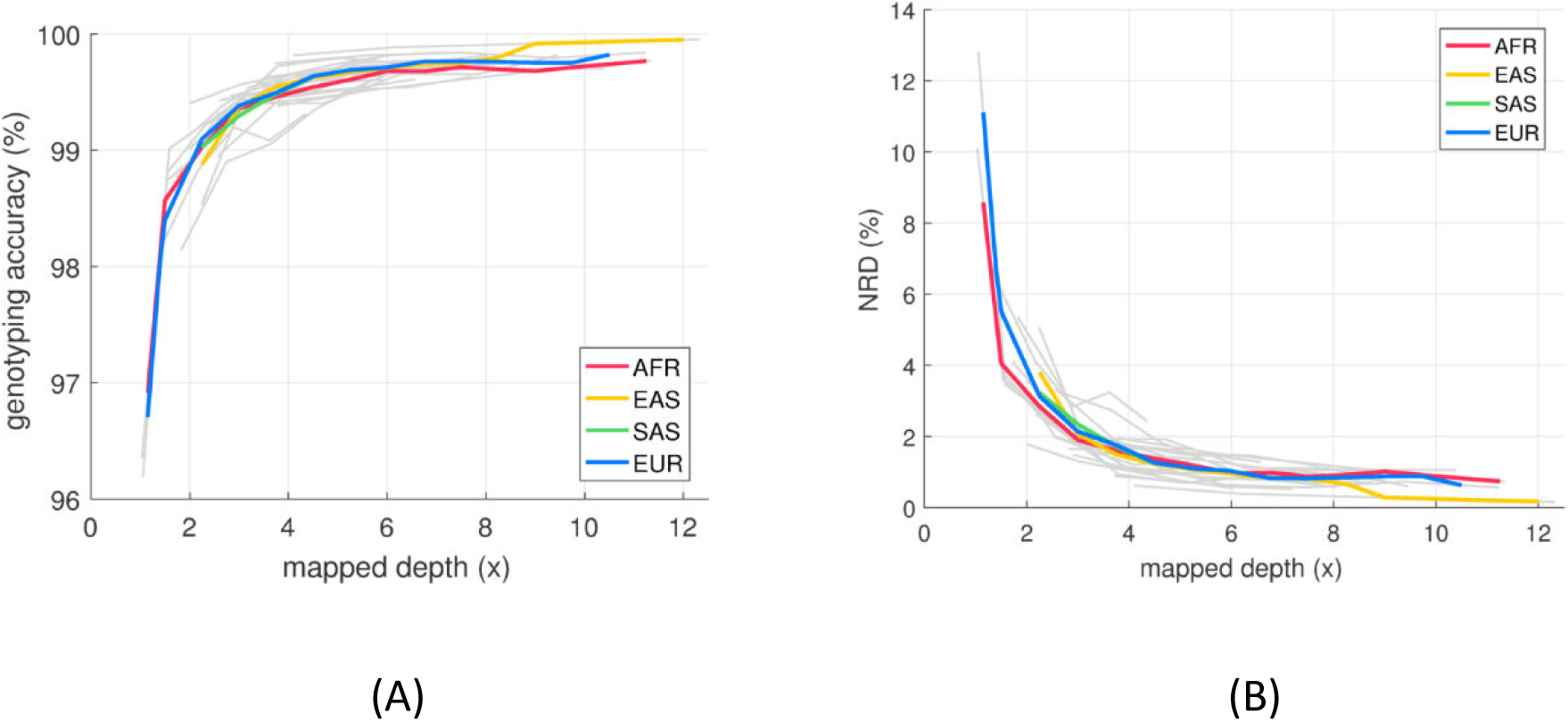
Coverage level affects genotyping performance. We applied the pipeline for analyzing very small datasets to three datasets separately; each dataset contains ten query samples randomly selected from the 1000 Genomes Project Phase 3. The BAM files of these query samples were downsampled to 100%, 83.3%, 66.7%, 50%, and 33.3% of the original coverage level. We measured the genotyping accuracy and non-reference discordance of every query sample at these coverage levels. The grey curves show the performance of each sample as a function of mapped depth. The colored curves show the aggregated performance of the samples from the same continent.

**Table 4.** Performance of applying the Ref-Reveel pipeline to a very small dataset with merely ten samples from unknown populations.

| query sample |  |  | top 5 populations |  | evaluated using Omni data |  |  |  |  | evaluated using CG data |  |  |
| --- | --- | --- | --- | --- | --- | --- | --- | --- | --- | --- | --- | --- |
| identifier | population<br>/ continent | depth<br>(x) | predicted according to<br>*F*-statistics |  | accuracy<br>(%) | NRD<br>(%) | RRD<br>(%) | RAD<br>(%) | AAD<br>(%) | het-accuracy<br>(%) | SN (%) | PR (%) |
| HG01985 | ACB/AFR | 6.04 | ASW,ACB,LWK,YRI,ESN |  | 99.732 | 0.753 | 0.189 | 0.336 | 0.535 | 98.665 | 99.051 | 92.135 |
| HG00759 | CDX/EAS | 8.51 | CDX,KHV,CHS,JPT,CHB |  | 99.691 | 0.666 | 0.192 | 0.436 | 0.463 | 98.497 | 98.971 | 91.465 |
| NA18525 | CHB/EAS | 12.33 | KHV,CHB,CHS,CDX,JPT |  | 99.931 | 0.160 | 0.000 | 0.000 | 0.490 | 98.505 | 98.978 | 91.462 |
| HG00171 | FIN/EUR | 3.19 | CLM,PUR,GBR,CEU,TSI |  | 99.541 | 1.458 | 0.260 | 0.832 | 0.986 | 98.929 | 99.247 | 92.394 |
| HG00096 | GBR/EUR | 4.66 | IBS,PUR,FIN,TSI,CEU |  | 99.628 | 1.026 | 0.291 | 0.440 | 0.671 | 98.913 | 99.229 | 92.417 |
| NA21092 | GIH/SAS | 5.17 | BEB,GIH,ITU,PJL,STU |  | 99.533 | 1.174 | 0.258 | 0.435 | 1.700 | 98.912 | 99.225 | 91.587 |
| NA18992 | JPT/EAS | 7.19 | CHB,CHS,JPT,KHV,CDX |  | 99.829 | 0.380 | 0.125 | 0.116 | 0.441 | 98.492 | 98.971 | 91.463 |
| HG01595 | KHV/EAS | 4.87 | KHV,CHB,JPT,CHS,CDX |  | 99.417 | 1.411 | 0.350 | 0.494 | 1.800 | 98.492 | 98.973 | 91.474 |
| NA19430 | LWK/AFR | 3.08 | ASW,LWK,ACB,YRI,ESN |  | 98.829 | 3.141 | 0.695 | 1.540 | 2.799 | 98.673 | 99.057 | 92.137 |
| NA20502 | TSI/EUR | 4.35 | IBS,TSI,FIN,CEU,PUR |  | 99.362 | 1.954 | 0.236 | 1.625 | 1.003 | 98.912 | 99.232 | 92.419 |

We also evaluated the genotyping performance of the samples in the combined VCF files, using the CG data as benchmarks. In every combined dataset, the number of samples that existed in the CG dataset as well ranged from 32 to 65. We measured the genotyping accuracy at the sites where the CG data reported heterozygous and homozygous non-reference. The measurement reflected the performance not only at common polymorphic sites, but also at rare variants sites, as the CG data was sequenced at a sufficiently high coverage. We further measured the sensitivity and precision, and reported the results in **Table 4**.

## Discussion

### Scalability

Whole-genome sequencing of large cohorts continues to become a promising trend. As such, developing an accurate genotype-calling method that is applicable at high scale, and practical for a very large number of individuals (for example, 1,000,000 individuals), becomes critical. In this paper, we introduced a novel method for large-scale variant calling, and demonstrated its applicability on one of the largest publicly available datasets, the UK10K data, which contains 3,910 whole genomes. When applying our method to a million whole genomes, a main bottleneck will be the pairwise LD estimation; the time complexity of this step is *O*(*m*^2^*n*). However, when the sample size scales up to such a large number, our method does not require the entire dataset for LD estimation. Rather, a subset of the individuals (such as, a few thousand) can be used for the estimation. By doing so, even if the sample size increases dramatically, the running time for LD estimation remains unchanged. Another bottleneck will be building simplified LD structures (see **Feature selection** of **Online Methods**). Like the first bottleneck, given a very large sample size, this step can be conducted from a subset of samples; moreover, the simplified LD structures built from a cohort can be reused for similar cohorts. Our experiments on the 3,910 UK10K samples shows that 32.65% and 50.60% of the total running time was spent on LD estimation and building simplified LD structures. The rest of the computation consumed 16.56% of the total running time; I/O costed only 0.19%. When the sample size scales up from 3,910 to one million, by downsampling the dataset to 3,910 individuals for the two bottleneck steps, the overhead of those two steps will remain the same. The rest of the overhead scales linearly with the sample size. Thus, the total computation cost will increase by 43 times.

### Guidelines for the generation of reference panels

Using a large reference panel has the potential to enhance the genotyping performance. **Supplementary Figure S2** shows that when the reference panel size was increased to 5,000, the genotyping accuracy was nearly saturated. With recent large-scale sequencing efforts, reference panels of comparable size are becoming available for a growing number of populations.

Ideally, the reference panel is composed of samples from the same population, or ancestral group, as the query samples. When a single-origin panel is unavailable, a panel that represents a more diverse set of ancestral population can be appropriate. **Supplementary Figure S3** shows that using the heterogeneous panel can improve the genotyping accuracy. Nevertheless, as expected, the effectiveness of using the heterogeneous panel was lower than when a single-origin panel was used.

**Supplementary Table S2** shows that even though the reference panel contains the genotypes at common polymorphisms only, incorporating such a panel into the genotyping process can reduce the error rate considerably. If the reference panel contains the genotype at all polymorphic sites, roughly two times of errors can be corrected, compared to the reference panel with common variants only.

### Coverage level of future sequencing projects

Reference-based genotype calling has the potential to further reduce the sequencing coverage needed for achieving high-accuracy genotyping. **Supplementary Figure S4** shows the performance of our method using a reference dataset. When utilizing a reference panel with a similar size to the UK10K Cohorts, and applying our method to a newly sequenced cohort of 2,500 individuals, a coverage level of roughly 2.5x is required to achieve a genotyping accuracy of 99.5%. Thus, with a lower average-coverage requirement, large-scale study efforts can afford the investigation of an even more comprehensive cohort.

## Acknowledgements

TwinsUK is funded by the Wellcome Trust, Medical Research Council, European Union, the National Institute for Health Research (NIHR)-funded BioResource, Clinical Research Facility and Biomedical Research Centre based at Guy’s and St Thomas’ NHS Foundation Trust in partnership with King’s College London.

## Authors’ contributions

LH and SB conceived the study and planned the experiments. LH designed the algorithm. LH and PD implemented the software and performed the experiments. LH, SB, and SB contributed to the manuscript. All authors read and approved the final manuscript.

## Online Methods

### Overview of Ref-Reveel

Ref-Reveel infers the genotypes for a cohort of *n* sequenced query individuals given a background cohort of *r* reference individuals. We assume that prior to the analysis, the genotypes of the reference individuals are known. Ref-Reveel discovers *m* likely polymorphic sites from the query data using the SNP-discovery method outlined in our previous work (Huang et al. 2016) and sequentially calls the genotypes at those sites.

Ref-Reveel is a multi-class AdaBoost based genotype caller. A weak learner infers the genotypes at all the markers across query genomes simultaneously. Building weak learners for every marker individually is not feasible, because the evidence at a single marker is often not sufficient for genotype calling due to low sequencing coverage. Our method creates a set of candidate weak learners before perform performing boosting. Those weak learners differ at the simplified LD structures they incorporate. After the weak learners are created, our multi-class AdaBoost algorithm is applied to every marker independently, using the reference panel as training data. The genotypes belong to three classes: homozygous reference (*g*=0), heterozygous (*g*=1), and homozygous alternate (*g*=2). The sample weight distribution is initialized as uniform across all the training samples. In every iteration, we pick the weak learner that minimizes the weighted sum of classification errors from constructed candidate weak learners, compute the weight of the picked weak learner, and update the sample weight distribution. We control overfitting by using *k*-fold cross validation (by default, *k*=10).

A weak learner, or a weak genotype caller, employs an iterative method (**Supplementary Figure S5**). In every iteration, our “guess” on the genotypes at all the markers are updated in a synchronized manner, that is, we refresh the genotypes only when the genotype inference from previous iteration for all the markers is ready. Ideally, we infer the genotype at a marker utilizing the comprehensive LD structure and the current estimation of the genotypes at all the markers. This strategy, obviously, is very time consuming. Instead, we build a local, simplified LD structure for every marker. A structure consists of several markers that have strong LD with the evaluated marker. Although these markers tend to locate near the evaluated marker, our structure does not include all the markers in the neighborhood. Instead, Ref-Reveel picks most informative markers based on LD, regardless of genetic distance. The strength of LD is evaluated on the fly from the input sequencing data of query genomes and, if applicable, the reference panel. The candidate weak learners differ at the metrics that are used to estimate the strength of LD, and consequently, they differ at the simplified LD structures.

A weak learner conducts multiple rounds of the iterative method described above. Each round starts with (re-)estimating pairwise LD and (re-)building simplified LD structures, using the output genotypes of the previous round as initial genotypes. The number of rounds can be specified by users. In our experiments, we apply three rounds because we have found that the LD estimation stabilizes within three rounds.

### Input data

Ref-Reveel accepts two sets of input information, regarding query genomes and reference genomes. For the newly sequenced query genomes, many large-scale projects provide genotype likelihoods instead of the raw sequencing data or alignments to avoid intensive input data volume and/or intensive data transfer. Besides relatively smaller data size, genotype likelihoods inherently embed the mapping quality of sequencing reads. Ref-Reveel accepts the genotype likelihoods of query genomes as inputs, and then calculates the *effective* reference and alternate read counts for internal use. Let *c* and *d* be the number of reads supporting reference and alternate alleles at the evaluated marker. We precompute a table of Pr{*c*, *d*|*g*} for every possible combination of reference and alternate counts *c*, *d* given an estimated error rate using the revised MAQ model (Li, 2010). For every target site, we find a combination of 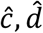 that minimizes the Manhattan distance between 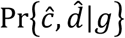 and the input genotype likelihoods at the target site over all the *g*’s. When the previously inferred genotypes of reference genomes are available, Ref-Reveel accepts those genotypes as additional inputs.

### Metrics for LD estimation

Pairwise LD estimation needs to be computational inexpensive because of the quadratic computation overhead with respect to the number of markers. For this reason, we define a family of metrics that can easily calculated from the effective read counts 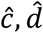; the conventional metrics, such as correlation coefficient, are more computationally expensive due to the need of preliminary genotype inference and phasing.

Let *X*_*i*,*s*_ be the event that at least one read at marker *i* of sample s supporting alternate allele, that is, 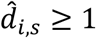. We categorize samples using the following relation predicates: *X*_*i*,*s*_ ∧ *X*_*j*,*s*_, ¬*X*_*i*,*s*_ ∧ *X*_*j*,*s*_, *X*_*i*,*s*_ ∧ ¬*X*_*j*,*s*_, and ¬*X*_*i*,*s*_ ∧ ¬*X*_*j*,*s*_. The samples with 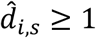 and 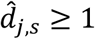, that is *X*_*i*,*s*_ ∧ *X*_*j*,*s*_, are the evidence of LD; we therefore use the number of such samples as the numerator of our metric. A naïve denominator is the total number of samples. This number, however, can easily be dominated by the number of samples with 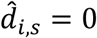 and 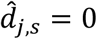, that is ¬*X*_*i*,*s*_ ∧ ¬*X*_*j*,*s*_, especially when markers *i* and *j* are rare variants. We therefore subtract this dominant number from the denominator. The metric can be written as follows:

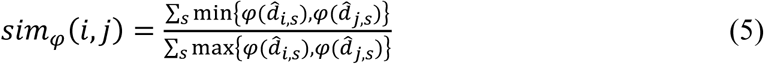

where, *φ*(*z*) = sgn(*z*). The sign function sgn(*x*) = 0 if *x* = 0, and sgn(*x*) = 1 if *x* > 0.

More broadly, this metric belongs to a family of metrics with different *φ*’s. We can easily define a series of metrics by picking different *φ* functions. In our experiment, we set *φ*(*z*) = *z*^*b*^ and vary the exponent *b* to create a set of candidate weak learners. Regardless of the choice of *φ* functions, the time complexity of *sim*_*φ*_ is linear to the number of samples.

The metric *sim*_*φ*_ can be calculated solely using the effective alternate read counts of query genomes or using the combination of query and reference data. The latter is recommended when the query and reference genomes follow similar LD patterns, for example, they come from the same population or ancestral group. The additional LD evidence provided by the reference genomes benefits the pairwise LD estimation especially when the number of query genomes is limited.

### Feature selection

To build a weak learner at an evaluated marker, we pick a collection of markers to form a simplified LD structure. The current estimated genotypes at those markers are referred to as the features of that weak learner. All the markers are candidates for feature selection. A naïve approach of feature selection is to pick a set of markers with highest *sim*_φ_ score. This filter feature selection method (Guyon and Elisseeff, 2003) is computationally efficient but only based on the correlation between the evaluated marker and other markers. The correlation between other markers is ignored. Thus, although each of these markers is one of the most informative features independently, the set may not be the most informative feature vector; when the chosen sites strongly correlate with each other, the information gained from selecting additional sites after the first one is limited.

To overcome this limit, our feature selection is a heuristic similar to Minimum Redundancy Maximum Relevance (Peng et al, 2005). To start with, all the markers are sorted based on their *sim*_*φ*_ score. Then, to construct a vector with *k* features, we conduct the following procedure *k* times (by default, *k* =3-5, depending on *n* + *r*): every time the marker with the highest feature score is added into the feature vector. Let *t* be the evaluated marker, and *M* be the set of markers in the current simplified LD structure, the feature score of marker *i* with respect to the evaluated marker *t* is defined as

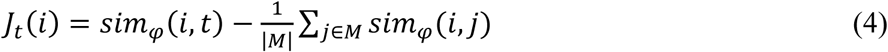

Different from Minimum Redundancy Maximum Relevance, which formulates *j*_*t*_(*i*) as a function of the information gain (Shannon, 2001), we calculate *j*_*t*_(*i*) using the *sim*_*φ*_ score to overcome the domination effect of ¬*X*_*i*,*s*_ ∧ ¬*X*_*j*,*s*_ cases.

### Weak learner

Our weak learners are implemented using a two-step iterative method (Huang et al. 2016), which is performed on all the markers across all the query genomes simultaneously. In the first step, given the current estimation of genotypes **G**, our method calculates the genotype probabilities **P**. As the samples are sequenced at a low coverage, the mapped reads have limited power to estimate the genotype probabilities **P** with high confidence. We therefore utilize the non-random association between alleles at different locations, that is, the linkage disequilibrium, to leverage evidence from additional sites in the calculation of a genotype probability. While the amount of evidence provided by the mapped reads for marker *i* is limited, when allele *A* and marker *i* and allele *B* at marker *j* have high levels of co-occurrence, observing allele *B* implies an increased chance of observing *A*. In the second step, the method infers the genotypes **G** by maximizing the current estimation of genotype probabilities; the predicted genotypes are then used to refine the genotype probabilities in the following iteration. We update **P** and **G** in a synchronous manner, that is, we first calculate the new values for **P** for every marker in every sample, and then overwrite all the old values with the new values before updating **G**. The two steps are iterated until the genotypes call converge.

Formally, we refer to a marker that is evaluated in a query sample as a target site and define the estimated genotype at the target site as *g*_*t*,*s*_, where *t* represents the evaluated marker and *s* is the evaluated query sample. Let *M* be the set of markers in the simplified LD structure of the evaluated marker. The estimated genotypes at the markers in set *M* in the evaluated sample *s* is denoted as vector **g**_*M*,s_. In the lth iteration, we estimate the genotype probability 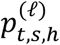 by calculating the probability that the target site exhibits a certain genotype *h* given **g**_*M*,s_ in the (l - 1) th iteration and read alignments at the target site, that is 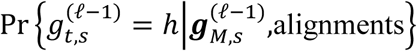, where *h* can be {0,1,2}, representing homozygous reference, heterozygous, homozygous alternate, respectively. Using the Bayes’ rule, we compute this conditional probability as follows, in which we do not need to calculate the denominator as it is identical to all the *h*’s:

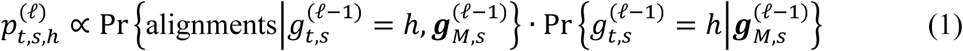

The first term calculates the probability of observing the alignments at the target given the genotypes at the target and the genotypes at the markers in set *M* in the evaluated sample. This term can be simplified as 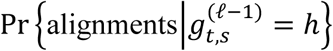 because of conditional independence. The probability Pr{alignments|*g*_*t,s*_ = *h*} is essentially the genotype likelihoods at the target site, which is pre-calculated from sequencing read alignments of query genomes. The conditional probability in the second term is identical across all the samples in the cohort. This is important because we can divide the computation of genotype probability of all the samples at a certain marker into two *O*(*n*) computations. First, we calculate 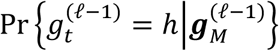 for *h* = {0,1,2} by accessing every sample once, where *s* is removed from the notation because this conditional probability aggregates across all the samples, including query samples and, if applicable, reference samples. Second, we calculate 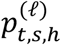 by simply retrieving the 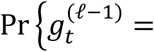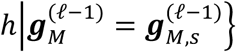 value and the genotype likelihood value. We simplify Equation (1) as

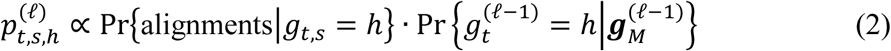

In the second step of our iterative method, the genotype probabilities for all the h’s are used to update the genotype:

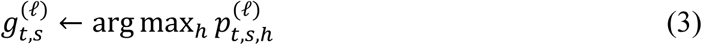

Our experiments on simulated data show that, the vast majority of loci converge within ten iterations; applying additional iterations yields little improvement in genotyping accuracy.

### Multi-class AdaBoost

Our multi-class AdaBoost-based algorithm starts with creating a set of candidate weak learners. A weak learner simultaneously infers the genotypes of query genomes at all the markers. The weak learner that is specified with LD metric function *φ* is referred to as *H*_*φ*_. To build *H*_*φ*_, we estimate pairwise LD by computing the metric *sim*_*φ*_ from the genotype likelihoods of query genomes and the genotypes of reference genomes. The resulting LD estimation is then used to build simplified LD structures for all the markers, referred to as ***M***_*φ*_. Using these simplified LD structures, applying our two-step iterative method to the genotype likelihoods of query genomes gives the weak learner *H*_*φ*_, yielding a genotype matrix ***G***_*φ*_.

We then calculate *H*_*φ*_(***x***_*i*_) (*i* = 1,…,*r*) for all reference genomes at all the markers simultaneously, where ***x***_*i*_ represents all the known information regarding the *i* th reference genome. In this process, we intentionally omit the LD estimation and the feature selection steps, and reuse the simplified LD structures ***M***_*φ*_. Based on ***M***_*φ*_, our two-step iterative method is applied to the reference genomes, assuming uniform genotyping likelihoods. The term 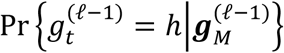 in Equation (2) is calculated using the genotypes of the reference genomes.

The next procedure is applied to every marker *t* separately, and formulated as follows:

- Initialize the set of chosen weak learners Ψ ← Ø.
- Repeat until cross validation indicates overfitting or all the candidate weak learners are chosen.
  - Step 1. Initialize uniform weight *w*_*i*_ (*i* = 1,…,*r*) over *r* labelled training samples (***x***_1_, *y*_1_),…,(***x***_*r*_, *y*_*r*_), where the class label *y*_*i*_ is the genotype of the ith reference genome at marker *t*.
  - Step 2. Choose the weak learner *H*_*φ*_* that minimizes the sum of the weighted classification errors 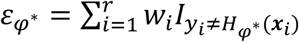, where *I*_*A*_ is an indicator function that equals one if A is true and zero otherwise.
  - Step 3. Calculate 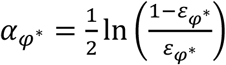.
  - Step 4. Update 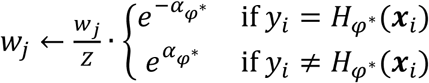, where Z is a normalization factor.
  - Step 5. *k*-fold Cross validation, starting with randomly partitioning the r training samples into *k* groups. A single group is retained as validation samples and (*k* - 1) groups are used as training samples. Then, our method applies Step 1-4 to the training (*k* - 1) groups, and then applies the final hypothesis at current stage to the validation samples. This process is repeated *k* times, rotating the validation group. The total validation error is accumulated over the *k*-fold cross validation.
  - Step 6. Add *φ*^*^ into *Ψ* if pass the *k*-fold cross validation.
- Apply the final hypothesis *H*_final_ = arg max_*y*∈{0,1,2}_ Σ_*φ*∈Ψ:*φφ*(***x***)_=*y* α_*φ*_ genomes, yielding final genotypes at marker *t*. to the query

Different from conventional AdaBoost-based algorithms (Freund and Schapire, 1997), Ref-Reveel builds weak learners and calculates *H*_*φ*_(***x***_*i*_) across all the markers before performing boosting. This is because utilizing LD structure is crucial in our two-step iterative genotype calling method. Training weak learners for every single marker individually is significantly less effective.

Recall that we vary the exponent *b* of *φ*(*z*) = *z*^*b*^ in our experiment to create a set of candidate weak learners. **Supplementary Figure S6** shows that the weak learner with large *b* value provides high genotyping accuracy at the markers with low MAF, while the weak learner with small *b* value is beneficial for the markers with high MAF. The figure also shows the most promising range of *b* value is between 0 and 3, except for the variants with AF < 0.1%. For those very rare variants, the highest genotyping accuracy is achieved at *b* = 4.75.

### Filters for quality control

To limit the false positives (defined in **Table 1**) resulting from the genotype calling process, we measure two ratios at every marker. Let *θ*_*X*_ be the probability that allele *X* exists at a marker in a sample. The function *f*(*θ*_*X*_) = *aθ*_*X*_/(1 + *a* - *θ*_*X*_) with *a* = 5 × 10^-6^ introduced in (Huang et al, 2016) is a monotonically increasing function with *f*(0) = 0 and *f*(1) = 1; its derivative is also monotonically increasing. This function has an important property that *f*(*θ*_*X*_) approaches 1 if and only if *θ*_*X*_ is very close to 1. Let score_*X*_ be the summation of *f*(*θ*_*X*_) over all the samples at the marker. We measure the ratio of score_*X*_ between major and minor alleles, and the ratio of allele frequencies between major and minor alleles. Because of the property of *f*(*θ*_*X*_), score_*X*_ can be used to approximate the number of samples in which we observe strong evidence for the existence of allele *X*. Thus, the two ratios described above should not be significant different. If at a marker these two ratios are highly inconsistent, we are not confident regarding the genotypes at that marker. That marker is not reported as polymorphic in the outputs.

The false positive rate can be further reduced by applying several hard filters. The following markers are also removed from the output polymorphic sites: (1) the markers that fail the Hardy-Weinberg equilibrium, using Pearson’s chi-squared test; (2) the sites that are called as common variants by our algorithm but not reported in the integrated call set of the 1000GP Phase 3; (3) the sites at which SAMtools calls multiple complex variants.

### Pipeline for analyzing very small datasets

To infer the genotype of a dataset with very limited number of individuals, we propose a Ref-Reveel pipeline as follows. We first apply Ref-Reveel to query samples using a reference panel. The purpose of this step was to obtain a rough estimate of genotypes of each query sample, which was not necessarily accurate but serves as an approximation to measure genetic diversity between that query sample and a cohort. Next, for each query sample we infer from which cohort the sample is originated. To be specific, we estimated the *F*-statistics between that sample and each possible cohort, and predict the top cohorts according to the *F*-statistics. We then generate a new VCF file by combining the query sample with the samples from chosen cohorts. Finally, we apply Ref-Reveel to the new VCF file without using any reference panels to obtain the final genotype calls for the query samples.

### Implementation of Ref-Reveel

The Ref-Reveel software was implemented in C. The software is publicly and freely available at http://reveel.stanford.edu. The program accepts two compressed/uncompressed VCF/BCF files as inputs: a reference panel with genotypes (GT field) and a query panel with genotype likelihoods (PL or GL field). For the reference panel, genotypes can be phased or unphased. These two files are simultaneously parsed using HTSlib, a C library for high-throughput sequencing data parsing.

To support the parallel execution of the genotype calling, the software provides a command that partitions the query panel into a series of consecutive non-overlapped genomic segments. The genotypes of each partition are inferred independently. Partitioning has little impact on the genotyping accuracy as long as the segment size is reasonable, as the majority of inter-marker LD extends to a few hundred kilobases at most (Reich et al., 2001; Schaffner et al., 2005). Two segmentation alternatives were implemented: users can either specify a desired segment length of a chunk (by default, 500 kb) or specify a desired number of markers in a chunk (by default, 12,000). The latter option is recommended, as it avoids cases in which too few markers are present in an evaluated segment. Such a scenario could diminish the effectiveness of genotype calling. Random access to VCF files is supported to accelerate the data parsing process.

